# Automated Integration of Multi-Slice Spatial Transcriptomics Data in 2D and 3D

**DOI:** 10.1101/2023.03.31.535025

**Authors:** Denis Bienroth, Natalie Charitakis, Dillon Wong, Sabrina Jaeger-Honz, Dimitar Garkov, Kevin I. Watt, Julian Stolper, Hazel Chambers-Smith, Duncan MacGregor, Bronwyn Christiansen, Adam T. Piers, Enzo R. Porrello, David A. Elliott, Karsten Klein, Hieu T. Nim, Falk Schreiber, Mirana Ramialison

**Affiliations:** Novo Nordisk Foundation Center for Stem Cell Medicine, Murdoch Children’s Research Institute, Parkville, Victoria, 3052, Australia; Department of Paediatrics, Faculty of Medicine, Dentistry & Health Sciences, University of Melbourne, Melbourne 3010, Australia; School of Computing and Information Systems, Faculty of Engineering and Information Technology, University of Melbourne, Parkville; Department of Computer and Information Science, University of Konstanz, Konstanz, 78464, Germany; Anatomical Pathology, The Royal Children’s Hospital, Melbourne, Victoria, 3052, Australia; Melbourne Centre for Cardiovascular Genomics and Regenerative Medicine, The Royal Children’s Hospital, Melbourne, Victoria, 3052, Australia; Department of Anatomy and Physiology, University of Melbourne, Parkville, Victoria, 3052, Australia; Australian Regenerative Medicine Institute, System Biology Institute, Monash University, Clayton, Melbourne, Victoria, 3168, Australia; Faculty of Information Technologies, Monash University, Melbourne, Victoria, 3168, Australia

**Author notes:** co-first authors (Denis Bienroth, Natalie Charitakis). co-senior authors (Enzo R. Porrello, David A. Elliott, Karsten Klein, Hieu T. Nim). co-corresponding authors Falk Schreiber Mirana Ramialison.

## Abstract

The field of spatial transcriptomics is rapidly evolving, with increasing sample complexity, resolution, and tissue size. Yet the field lacks comprehensive solutions for automated integration and analysis of multi-slice data in either stacked (3D) or co-planar (2D) formation. To address this, we developed VR-Omics, a free, platform-agnostic software that distinctively provides end-to-end automated processing of multi-slice data through a biologist-friendly interface. Benchmarking against existing methods demonstrates VR-Omics’ unique strengths to perform comprehensive end-to-end analysis of multi-slice stacked data. Applied to rare paediatric cardiac rhabdomyomas, VR-Omics uncovered previously undetected dysregulated metabolic networks through co-planar slice analysis, demonstrating its potential for biological discoveries.

## BACKGROUND

Spatial transcriptomics (ST) is a rapidly evolving technology allowing the capture of spatial gene expression at unprecedented resolution to advance our comprehension of cellular processes and molecular mechanisms [1, 2]. Since many tissue samples being profiled exceed a standard vendor-provided chip size, analysing multi-slice ST data is an important task that cannot be fully supported using current single-slice tools [3]. Multi-slice samples are typically co-planar, where slices lie on the same plane (*e.g.* twelve prostate cryosections sections [4]) or 3D stacked, where parallel slices are orthogonally adjacent (*e.g*. thirteen E16.5 mouse embryo sections [5]). Bridging this technical gap enables tissues to be studied in their complete native spatial context, providing crucial insights, and understanding of the tissue structure and characteristics.

Several ST algorithms have been developed to process and mine ST multi-slice data, including Stitch3D [6] and VT3D [7]. However, these specialised multi-slice pipelines or analytical programmes handle only a narrow scope within the end-to-end analysis pipeline while also requiring advanced computational skills, creating a substantial barrier for end-users. Several online portals allow viewing of existing multi-slice ST datasets (SpatialDB [8], STOmicsDB [9], SODB [10], SOAR [11]) but limit the user to the datasets that are publicly available and do not allow analyses of user-uploaded multi-slice data. Finally, no current multi-slice ST tools allow manual interactive alignment of multi-slice data, both co-planar and 3D stacked formations, providing an interactive graphical user interface.

Here we present VR-Omics, an interactive, end-to-end platform to process, analyse and visualise ST data, all through a user-friendly graphical user interface (GUI). VR-Omics is designed to democratize ST data analysis, making it accessible to both bioinformaticians and non-bioinformaticians while generating reproducible results in a timesaving manner. As inputs, VR-Omics supports data from both sequencing and imaging-based ST technologies, as well as custom ST data matrices [12–14]. VR-Omics primarily functions as a conventional desktop application, with the inclusion of VR integration as an optional supplementary feature. While fully supporting single-slice ST data, VR-Omics further facilitates multi-slice ST data with additional capabilities, including merging or manual alignment of slides and comprehensive 3D visualisation.

We demonstrate the utility of VR-Omics through the analysis of both single and multi-slice rare paediatric cardiac rhabdomyomas (cRMs) for which the disease mechanism remains unknown [15, 16]. Using VR-omics, we identified novel transcriptional signatures distributed in discrete niches across the cRM. Additionally, we showcased the precision of VR-Omics by performing a meticulous three-dimensional (3D) reconstruction of cardiac structures, rigorously validating its performance against state-of-the-art ST tools. These multifaceted use-cases collectively exemplify the versatility and robustness of VR-Omics as a pivotal tool for effective and novel spatial transcriptomics research.

## RESULTS

### VR-Omics enables end-to-end ST analyses in 2D and 3D

With VR-Omics, we developed the first automated interactive tool with a graphical user interface (GUI) that allows both the concatenation and analysis of multiple co-planar slices, as well as the generation of a harmonised coordinate system of stacked multi-slice tissues. This integration is achieved through the utilisation of the Unity Gaming Engine combined with Python-based analysis workflows (**Supplementary Methods**).

For the analysis of multiple co-planar slides, we have engineered a user interface utilising the Unity engine (**Fig. 1a**). This interface enables users to visualise and manipulate tissue slides, allowing them to be freely arranged and rotated to approximate the original structure of the tissue. This spatial arrangement is captured in the *world space* of Unity, where the relative positions of the tissue slides are meticulously collected. These coordinates are subsequently transferred to a Python-based workflow, which concatenates the data objects according to their spatial alignment. Rotation of the slices is facilitated by applying transformation matrices to the array of data points. Once aligned, the separate data objects are concatenated into a single entity and converted into required formats typically compatible with downstream processes. Additionally, the interface provides options for users to input filtering and processing parameters necessary for subsequent analysis.

**Figure 1.**
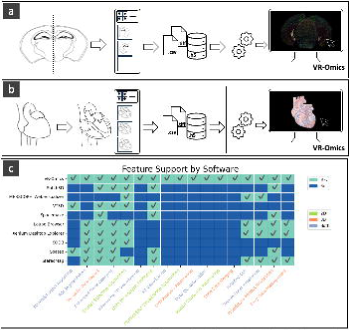
Analytical capabilities of VR-Omics. **a**, Schematic overview of data generation and processing workflow for multi-slice co-planar ST datasets. **b**, Schematic overview of data generation and processing workflow for multi-slice 3D stacked ST datasets. **c**, Heatmap comparison of analytical features supported by VR-Omics and other available software for ST analysis. Software is outlined on the y-axis while analytical features are listed on the x-axis and coloured according to their applicability to multi-slice co-planar or 3D stacked datasets. Features supported by the software are displayed in teal while unsupported features are displayed in dark blue.

In a parallel development for native 3D structures, VR-Omics includes a similar Unity-based user interface that supports the rotation and alignment of consecutive sections (**Fig. 1b**). This functionality allows for the visualisation of these sections as a cohesive 3D dataset, providing a comprehensive view of the tissue’s architecture.

These automated workflows, enhanced with a user-friendly interface, not only streamline the analysis of multi-sliced tissue sections but also minimise inconsistent analytical steps or parameters across slices. This is achieved by standardising the analysis process across various open-source programming languages, promoting consistency, and reducing variability that arises when using multiple packages and coding languages.

To demonstrate the comprehensive capabilities of VR-Omics, we conducted an extensive evaluation on common analysis workflows encompassing data input/export handling, visualisation, and general utility, among a broad spectrum of ST tools outlined in **Fig. 1c** [6, 7, 10, 13, 17–21]. We systematically identified the essential steps required for analysing both co-planar multi-sliced tissues and depth-stacked datasets. Our comparison with a diverse array of SRT tools revealed that VR-Omics is the only platform capable of executing all necessary procedures integral to this analysis (**Fig. 1c**). This underscores the necessity of a unified platform that consolidates comprehensive data analysis within a single environment, as these tasks often require integrating multiple tools and navigating diverse coding languages such as R and Python.

### 3D reconstruction workflow of the developing human heart

To illustrate the capabilities of VR-Omics in comparison against multi-slice ST tools, we performed an in-depth analysis of a 3D multi-sliced dataset derived from the developing human heart using publicly available data (**Fig 2, Supplementary Table 1**). Our comparative analysis employed a five-step workflow designed to cover all the necessary procedures for standard differentially expressed gene (DEG) analysis, applied specifically to spatial gene expression in different chambers of the developing human heart using a published 3D SRT dataset at 6 days post-conception (dpc) [22]. The workflow included: (1) inputting SRT data; (2) clustering spots/locations; (3) assembling slices into a 3D structure; (4) manually selecting two regions of interest (ROIs); and (5) identifying the top DEGs between these ROIs (**Fig. 2a**). This approach is critical for unraveling spatial gene expression patterns and comprehending the architectural complexities of tissues, where preserving the native 3D context is crucial for understanding the developmental intricacies that shape complex organ structures such as the heart.

**Figure 2.**
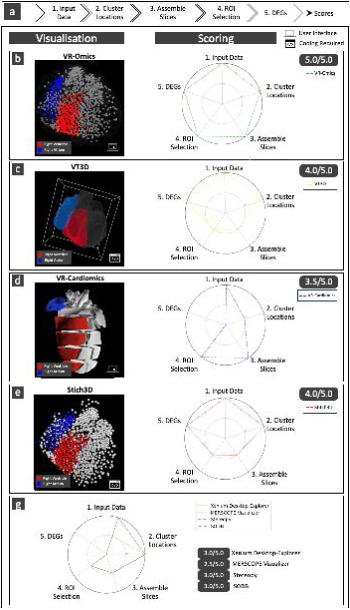
Cross-platform comparison for 3D reconstruction workflow of the developing human heart. **a,** Schematic of workflow to identify differentially expressed genes between the right atrium and right ventricle. **b**, Results from VR-Omics completing the workflow. On the left the final visualisation of the right atrium (blue) and ventricle (red) are displayed within the VR-Omics GUI and the spider plot on the right displays VR-Omics ability to complete all 5 steps of the workflow. **c**, Results from VT3D executing the workflow. On the left the final visualisation of the right atrium and ventricle is displayed within VT3D and the spider plot on the right displays its ability to complete 4/5 steps of the workflow. **d**, Results from VR-Cardiomics executing the workflow using a mouse heart, with the chambers displayed on the left and the spider plot on the right displaying the 3.5 steps that can be completed within VR-Cardiomics. **e**, Example results from Stich3D executing the workflow with chambers displayed and coloured on the left and the spider plot on the right. Selection of specific ROIs to highlight the right atrium in black and right ventricle in red performed in R outside of the STich3D framework. **g**, Spider plot demonstrating performance of remaining tools trailed for this workflow.

We conducted this workflow comparing VR-Omics against several other tools: STitch3D [23], VR-Cardiomics [12], Loupe Browser [19], Xenium Desktop-Explorer [18], MERSCOPE Web-Vizualizer [20], Stereopy [17], SODBView (through its visualiser SOView) [10] and VT3D [7] (**Fig. 2, Supplementary Figure 1, Supplementary Table 1, Supplementary Methods**). Each tool was evaluated based on its ability to complete each workflow step, with scores assigned for full (score=1), partial (score=0.5), or unsuccessful (score=0) execution.

Firstly, the comparative analysis underscored that most tools are tailored primarily for 2D data, with limited capabilities for integrating 2D slices into a coherent 3D structure. Accordingly, all tools compared except VR-Omics, VR-Cardiomics, Stitch3D and VT3D were not able to proceed beyond completing task 3. Secondly, while VT3D and STitch3D support the assembly of 2D slices into 3D, they lack functionality for native 3D ROI selection, limiting the ability to isolate and investigate spatially restricted cell populations across the 3D stack. Thirdly, our previous tool, VR-Cardiomics, facilitates steps 3 to 5 of the workflow but is constrained by its inability to process novel input data, highlighting VR-Omics’ unique flexibility and comprehensive support for 3D SRT data analysis [12]. Notably, only VR-Omics achieved a perfect score across all five steps, showcasing its unparalleled capacity for conducting comprehensive 3D spatial analyses and DEG identification (**Supplementary Tables 1-2**).

### Understanding molecular signatures of paediatric cardiac rhabdomyomas using VR-Omics

To further illustrate VR-Omics’ unique end-to-end capabilities in the analysis of multi-slice datasets, we tested it on an original co-planar dataset to interrogate a disease mechanism that has not previously been transcriptomically investigated. Cardiac rhabdomyomas (cRMs) are non-cancerous tumours that, while extremely rare in the general population, with a prevalence between 0.0017%-0.28%, are the most common form of primary cardiac paediatric tumour [16, 24]. Depending on the anatomical presentation of cRMs in cardiac muscle these tumours can impair normal heart function in a paediatric cohort by causing respiratory distress, arrythmias, ventricular obstructions and cardiac failure [25, 26]. When their presence leads to severe health complications, treatment options are very limited and include surgical resection [25], which may lead to additional mortality and morbidity complications for the patient [24]. Yet, the molecular mechanisms underlaying cRM formation remain unknown. Limited transcriptomics studies of 10 marker genes have been performed on these tumours, identifying abnormal regulation of the mTOR signalling pathway associated with cell proliferation [27]. To elucidate the molecular and cellular perturbations leading to these abnormal cell populations and to explore tumour heterogeneity in a spatial context, we profiled multiple cRMs with 10X Genomics Visium platforms and VR-Omics.

Samples were obtained from cRMs located in the ventricles of two separate patients. The first sections, MCHTB302, were obtained from a patient diagnosed with Birt-Hogg-Dubé Syndrome (BHDS) who had two cRMs, one in the left ventricular cavity and a second in the apex of the heart (**Fig. 3a_i_**). For this study, two sections were taken from the cRM located in the left ventricular cavity (**Fig. 3a_ii__-iii_**). Initial immunofluorescence staining performed on this tumour indicated evidence of proliferating cardiomyocytes through colocalized staining of cTNT and Ki-67 (**Fig. 3a_iv_**) [28]. The section originating from the center of the tumour was then divided across 3 Visium slides, due to size limitations of the Visium capture area (**Fig 3b-c**). The 3 slides were easily and quickly recombined using VR-Omics to generate a single data object from the multi-slice, co-planar dataset that more faithfully recapitulated the biological sample (**Fig. 3b-c**). Using the STAGATE algorithm which was specifically developed for SRT clustering and is implemented in VR-Omics’ AW, we identified cell populations (cluster 0 and 4) in this dataset that displayed a switch in cellular metabolism to increased aerobic respiration and oxidative phosphorylation when their markers (*ACO2, ATP5F1A, ATP5F1B, ATP5ME, COX4I1*) were used as an input for a gene ontology enrichment analysis (**Fig. 3d**). These populations may generate additional ATP through upregulation of genes such as *ATP5IFI,* paralleling aspects of the Warburg effect which sustains abnormal cell proliferation in cancerous tumours [29]. These populations may generate additional ATP through upregulation of genes such as *ATP5IFI,* paralleling aspects of the Warburg effect which sustains abnormal cell proliferation in cancerous tumours [29]. These findings are in line with metabolic changes observed in murine models with mutated or knocked out *FLCN*, similar to the patient’s genotype, that are proposed to be due to the dysregulation of the AMPK-mTOR pathway [30, 31]*. FLCN* has previously been suggested to play a role in cardiac homeostasis and this lack of regulated cellular energy levels may be driving this abnormal proliferation of cardiomyocytes [30]. *FLCN* is implicated in other cardiac diseases such as cardiac hypertrophy, and therefore may be one of many dysregulated genes giving rise to cRMs [30].

**Figure 3.**
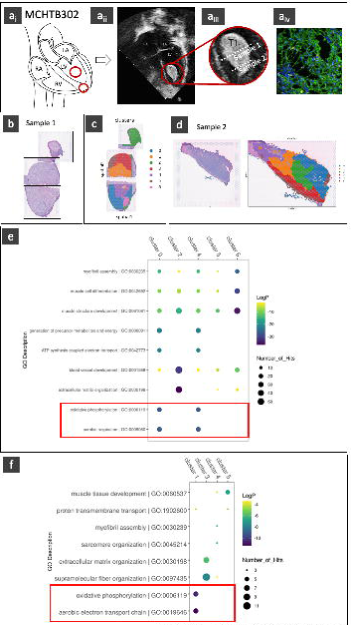
Analysis of multi-slice co-planar and single slice ventricular MCHTB302 cardiac rhabdomyoma data. **a_i_**, Schematic representing location of cRM used for data generation. **a_ii_**, ECG displaying location of cRM in left ventricular cavity. **a_iii_**, Schematic representing approximate locations of sections taken for data generation using the 10X Genomics Visium platform. **a_iv_,** Immunostaining with Ki67 indicating proliferation within the tumour (orange); nuclei staining (Hoescht, blue); cardiac troponin T (cTNT, green). **b,** Reconstruction and concatenation within VR-Omics of 3 co-planar, H&E stained slices separated onto different Visium capture areas due to size of the tissue. **c**, Example of STAGATE clusters visualised across the reconstructed co-planar dataset (labelled as Sample 1 in panel a) within VR-Omics. **d**, Example of STAGATE clusters visualised across the single section dataset (labelled as Sample 2 in panel a) within VR-Omics. **e**, GO analysis results of multi-slice coplanar dataset using the STAGATE cluster markers as inputs for Metascape. Oxidative phosphorylation and aerobic respiration signatures are highlighted in a red box for clusters 1 and 5 located at the centre of the tumour. **f**, GO analysis results of single-slice dataset using the STAGATE cluster markers as inputs for Metascape. Oxidative phosphorylation and aerobic electron transport chain signatures are highlighted in a red box for cluster 1 located at the centre of the tumour.

We then sought to validate this upregulation of oxidative phosphorylation and aerobic respiration in the clusters identified through the STAGATE algorithm. First, we validated the signature against an additional tumour section taken from the same patient, that did not need to be divided across Visium capture areas (**Fig. 3e**). This served to confirm that the molecular signatures were not an artefact from VR-Omics’ multi-slice, co-planar analytical approach. Using the same analytical pipeline this additional sample from MCHTB302 displayed the same signature for up-regulation of pathways related to aerobic respiration in clusters located in the centre of the tumour (**Fig. 3f**). Therefore, we demonstrate that VR-Omics’ provides a unique capacity for analysing co-planar datasets to explore novel biological mechanisms in rare tissue sections.

We were fortunate to have access to samples from a second patient to investigate the presence of the transcriptional signature across cRMs with distinct etiologies. The second patient MCHTB361 presented with a cRM in the left ventricular outflow tract and had been diagnosed with Tuberous Sclerosis Complex (TSC) which is commonly linked to cRMs [27] (**Fig 4a**). Three separate sections were taken from this cRM, and each was captured using individual Visium slides (**Fig. 4a**). Across all three sections, the same signatures of upregulation of aerobic respiration and oxidative phosphorylation were present in clusters in the centre of the tumour (**Fig. 4b_ii_ -c, Supplementary Fig. 9**). Within these predominantly abnormal cardiomyocyte clusters (**Fig. 3 & Fig. 4**), there was evidence for altered transcriptional signatures through increased expression of genes such as *SLC25A4* which is consistent with increased metabolic activity [31]. Certain cell clusters also displayed upregulation of genes involved in the increase of glycolysis, potentially indicating the presence of the Warburg effect which may contribute to the growth of these benign tumours [29]. Moreover, the clustering performed across samples reveals additional transcriptional heterogeneity that cannot be identified using histological analysis of single-slice data sets (**Fig. 2 & Fig. 3**).

**Figure 4.**
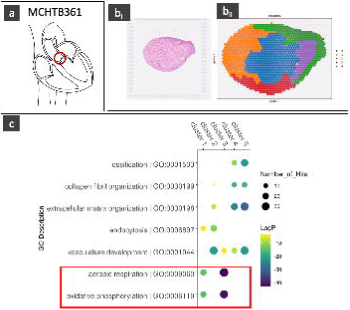
Analysis of single-slice MCHTB361 cardiac rhabdomyoma data. **a**, Schematic representing location of cRM used for data generation. **b,** Analysis of a single slice MCHTB361 dataset within VR-Omics. **b_i_**, H&E stained section visualised with VR-Omics**. b_ii_,** Example of STAGATE clusters visualised dataset within VR-Omics. **c**, GO analysis results of single slice dataset using the STAGATE cluster markers as inputs for Metascape. Oxidative phosphorylation and aerobic respiration signatures are highlighted in a red box for clusters 1 and 5 located at the centre of the tumour. **f**, GO analysis results of single-slice dataset using the STAGATE cluster markers as inputs for Metascape. Oxidative phosphorylation and aerobic electron transport chain signatures are highlighted in a red box for clusters 1 and 3.

These two avenues for validation demonstrate that the molecular signatures we identified are robust in both intra- and inter-patient samples. The ability to generate a single data object across multiple planar Visium sections allowed for a better understanding of the distribution of cell populations across the tumour, which would have otherwise been lost due to artificial tissue separation across capture areas. VR-Omics GUI makes this process easy and intuitive by simplifying data joining to a simple ‘drag and drop’ alignment exercise of Visium slices within the VR-Omic’s GUI, eliminating the need for computational steps to generate a single data object (**Supplementary Video 1**).

Subsequently we expanded our analysis to samples from a third patient (MCTHB362) who presented with two cRMs, in the left atrium and atrial valve, and whose father was a known carrier of TSC (**Fig. 4c**). The goal was to investigate further similarities between patients and identify potential differences that may arise due to atrial and ventricular signatures between the cRMs. For MCHTB362, tissue from both cRMs had to be sectioned to accommodate Visium size restrictions and were later virtually reconstructed like the samples from MCHTB302 (**Fig. 4c & Supplementary Fig. 8e**). This allowed direct comparison of multi-slice, co-planar datasets between patients to determine if similar signatures were present. Moreover, the presence of a single section that fit on an individual Visium slide (**Fig. 4c**) allows for intra-sample comparison for merged or individual data objects. In this instance, we noticed that the cRMs originating in the atria had a slightly different transcriptional profile than previous ventricular samples (**Fig. 3 & Fig. 4d**). However, upregulation of aerobic metabolism in certain clusters remained constant, demonstrating a consistent association with the tumour phenotype across all patients (**Fig. 3 & Fig. 4d**). Processing and visualization of multiple patient samples through VR-Omics facilitated the discovery of similar cell populations and molecular signatures in MCTHB302 and MCHTB361 that suggest they are characteristic of cRMs (**Fig 3-5 and Supplementary Fig. 9**). This signature was also present within MCHTB362 despite these samples being histologically and transcriptionally distinct from MCHTB302 and MCHTB361 (**Fig. 3-5 and Supplementary Fig. 9**). These findings across patients were further orthogonally validated outside of the VR-Omics framework (**Supplementary Results**). Altogether this provides evidence that this switch to an altered transcriptional signature to increase aerobic respiration is characteristic of the cRMs in all three patients (**Fig 3-5 and Supplementary Fig 9**).

**Figure 5.**
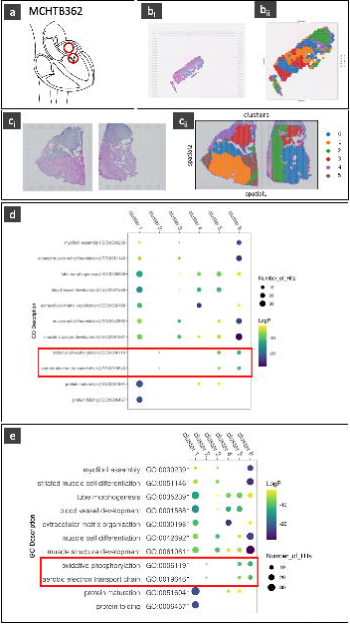
Analysis of multi-slice co-planar and single slice MCHTB362 atrial cardiac rhabdomyoma data. **a**, Schematic representing location of cRMs used for data generation using the 10X Genomics Visium platform. **b_i_**, H&E stained single slice section visualised with VR-Omics. **b_ii_**, Example of STAGATE clusters visualised across a single-slice dataset within VR-Omics. **c_i_,** Reconstruction and concatenation within VR-Omics of 2 co-planar, H&E stained slices separated onto different Visium capture areas due to size of the tissue. **c_ii_**, Example of STAGATE clusters visualised across the reconstructed co-planar dataset within VR-Omics. **d**, GO analysis results of single dataset using the STAGATE cluster markers as inputs for Metascape. Oxidative phosphorylation and aerobic electron transport chain signatures are highlighted in a red box for clusters 2, 5 and 6 located at the centre of the tumour. **e**, GO analysis results of multi-slice, co-planar dataset using the STAGATE cluster markers as inputs for Metascape. Oxidative phosphorylation and cellular respiration signatures are highlighted in a red box for clusters 2,3 and 6 located at the centre of the tumour.

## DISCUSSION

The number of multi-slice ST experiments is growing rapidly [32], simultaneously presenting a great opportunity for spatial biology investigations and a data science challenge. Addressing To address the gap in technologies capable of handling multi-slice data, VR-Omics was designed from the ground up to manage multi-slice ST data (both co-planar and 3D stacked) through an intuitive, customizable graphical interface that requires minimal computational skills (**Supplementary Fig. 2, Supplementary Video 1**). When compared to multiple tools and pipelines for SRT data analysis, VR-Omics demonstrates its unique capability in performing comprehensive and advanced end-to-end data analysis for multi-slice SRT datasets.

Benchmarking VR-Omics against most common SRT data analysis and visualisation tools using the 3D human developing heart dataset reveals that only VR-Omics fulfils all the necessary criteria for end-to-end analysis and visualisation typically of interest to biologists in a 3D stacked dataset.

The space of SRT data analysis urgently requires applications developed for 3D datasets to align, visualise and process multi-sliced data. At the time of writing, only STich3D [6], VT3D [7] and our previous work VR-Cardiomics [12] support 3D data visualisation and analyses besides VR-Omics. However, these applications focus on delivering specific aspects of the analytical pipeline, and therefore lack critical features to enable end-to-end analysis for users with a range of computational expertise. For example, VR-Cardiomics lacking the ability to analyse users’ novel data, rather focusing on visualisation of a singular pre-processed dataset. Which removes the flexibility of extending the tool to new studies. Thus, we have developed VR-Omics to bridge this gap, allowing users to analyse and interact with their 3D data in an intuitive manner.

Our comparative analysis between VR-Omics and other proprietary and open-source tools for ST multi-slice analysis (**Fig. 1c**), reflects a wide range of support for various SRT platforms and the advanced analytical tasks required by researchers. Despite the inclusion of tools specialised for enhancing spatial analysis in three dimensions, such as Stitch3D and VT3D, which primarily focus on visualization and data mining, only VR-Omics was able to complete all five essential steps for a comprehensive analysis (**Fig 2, Supplementary Table 1**). This evaluation confirmed VR-Omics’ unique ability to support a complete analytical workflow, significantly outperforming existing methodologies in efficiency and thoroughness of data analysis. VR-Omics’ singular success in completing all steps of the biological case study, particularly in identifying differentially expressed genes between two cardiac regions, further highlights its unparalleled capability to provide a robust and comprehensive framework for analysing complex spatial transcriptomics data.

Through the analysis of rare cRM samples we demonstrate that VR-Omics overcomes two major challenges in multi-slice co-planar ST data: (1) biologically adjacent samples being separated for processing which need to be merged for analysis, and (2) inter-sample comparisons. VR-Omics’ capability to analyse multi-slice datasets and easily perform inter-sample comparisons permitted the elucidation of key molecular mechanisms contributing to the growth of rare cardiac rhabdomyoma samples. The workflow available within the VR-Omics framework identified shared signatures of increased aerobic respiration across patients within abnormal cardiomyocytes faster than comparable workflows implemented across multiple disaggregated packages. This indication of potential dysregulation of the AMPK-mTOR pathway is known to have an effect on mitochondrial metabolism and has been demonstrated in murine models with mutations in *FLCN*, the same gene known to be mutated in patient MCHTB302 [30]. Without effective analysis of SRT data through framework tools such as VR-Omics, rare clinical samples preserved in FFPE such as these cRMs (prevalence 0.0017%-0.28%), would remain inaccessible.

## CONCLUSION

Multi-slice ST data analysis is quickly evolving [33], and VR-Omics rapidly evolves to address development and improvement cycles. VR-Omics natively supports multi-slice ST data from Visium, Xenium, Stereo-Seq and MERFISH, and will support Nanostring (CosMX) and Slide-Seq V2 in the near future. By addressing existing computational limitations in current multi-slice ST tools, VR-Omics offers unique advantages as demonstrated in this study using both stacked and co-planar multi-slice datasets. In this study we leverage VR-Omics to dissect the mechanisms underlying growth of paediatric cRMs. Our discovery unlocks potential insights into the regeneration of cardiac muscle for therapeutic purposes as the tumours develop within striated muscle and cause abnormal levels of proliferation after birth. VR-Omics stands out from other tools with unique software architecture, immersive environment, cross-platform compatibility, and a fully automated workflow, all aiming at simplifying and democratising ST analyses in the spatial biology community.

## MATERIAL AND METHODS

### VR-Omics Architecture

To deliver a seamless user experience, VR-Omics was conceived as a sophisticated end-to-end analysis tool, integrating two primary components via a user-friendly interface (**Supplementary Fig. 3**). The first component, the purpose-built Automated Workflow, focuses on the processing and mining of raw SRT data (**Supplementary Fig 3a, b**). The second component, the Visualiser, utilises the processed data to facilitate advanced data visualization, equipped with multiple features for enhanced data interaction and exploration through a fully interactive interface, designed with intuitive data mining processes in mind (**Supplementary Fig 3c, d**).

VR-Omics was developed utilising the Unity game development engine by Unity Technologies, starting with Unity version 2019.4.26f1. An upgrade to version 2021.3.11f1 (Long-Term Support, or LTS) was later implemented to ensure complete compatibility with the Virtual Reality toolkit plugin (XR Interaction Toolkit version 2.0.3) (**Supplementary Methods**). For each integrated SRT platform, we developed specific state-of-the-art Automated Workflows using Python executables and open-sourced tools and packages, which are embedded within the application to function as a unified software solution.

### Features of VR-Omics

#### Automated Workflow

For each of the integrated SRT platforms, a purpose-built Automated Workflow (AW) was developed to optimize data processing and allow advanced analysis of the SRT raw data. This optimization is achieved through a user-friendly interface that enables end-to-end processing and analysis of raw data, thus eliminating the need for time-consuming data formatting steps. All proprietary data is supported as provided by the vendor, and VR-Omics automatically detects the necessary files by selecting the main directory of each sample, reducing the risk of incorrect file selection. Within the AW, users can perform a variety of operations to tailor data analysis (**Supplementary Fig. 2a-d**), including standard filtering metrics to ensure data quality such as: (i) number of counts per location, (ii) percentage of overall mitochondrial gene expression, and (iii) number of cells expressing a certain gene. These metrics can all be easily visualised across the tissue section to highlight regions of poor data quality and capture inter- or intra-tissue heterogeneity (**Supplementary Fig. 2a**).

The default normalisation methods (as recommended by each analysis package) are applied, and data is log-transformed before highly variable genes (HVGs) are identified. Subsequently, principal component analysis (PCA) is performed, neighbours are computed, and UMAP or t-SNE coordinates are generated prior to performing spatially agnostic Leiden clustering or STAGATE-enhanced Leiden clustering depending on the dataset (**Supplementary Fig. 2b**). The AW executes spatial analyses using various Python libraries such as Scanpy, SpatialDE, STAGATE, and SpaceFlow, among others, for different commercialised and open-source SRT platforms and custom data (**Supplementary Fig. 3b**).

Spatially variable genes (SVGs), crucial for understanding cell differentiation and tissue organisation, are identified using SpatialDE for Visium data or Moran’s I statistic for other data types (**Supplementary Fig. 2c**) [34]. These steps, selected to follow a typical workflow of a bioinformatician, ensure VR-Omics provides a complete experience for the analysis. Additional capabilities for Visium datasets include the option to concatenate multiple sections from the same tissue in 2D for downstream joint analysis (**Supplementary Fig. 2e**) and the exploration of clustering results using the Leiden algorithm or STAGATE for more complex analyses.

#### Visualiser Capabilities

Across these SRT platforms, different terms are used for the smallest individual capture area of each technique, *e.g*., *spots*, *locus*/*loci*, *bins,* or *locations.* For consistency we will henceforth use the term *locations*, always referring to the smallest individual capture area of the respective SRT methods.

After processing their data through the Automated Workflow (AW) or loading the requisite files, irrespective of the data platform used, users can explore their datasets within the Visualiser. The Visualiser offers an array of interactive features designed with input from biologists to create intuitive analysis tools and ensure availability of the right resources to address various biological questions. These features support the systematic exploration and comparison of gene expression data across samples within a 2D desktop or VR environment (**Supplementary Fig. 4-5, Supplementary Table 3**).

Key capabilities include the visualisation of gene expression, enabling insights into the expression of significant markers, such as EPCAM in breast cancer, which is associated with poor prognosis [35]. This visualization aids in correlating disease severity with tissue characteristics, further enhanced by the superimposition of metadata like clustering results or images alongside gene expression data, allowing for rapid assessment of alignment between clustering, gene expression, and tissue histology.

The platform also enables the exploration of gene expression profiles, crucial for leveraging the full potential of SRT data. Features like visualising two different gene expression profiles on adjacent slides or within the same tissue section, as well as vector-based gene expression differences, facilitate comparisons of markers linked to disease prognosis or aggressiveness such as *EPCAM* and *ELF3* in breast cancer [35, 36]. This comparison is valuable for understanding their impact on specific regions or cells within a tumour. Replication of actions across duplicated slides, such as region of interest (ROI) selection, is supported.

Gene search functionality allows users to find and visualise a gene of interest’s expression through a heatmap, showcasing relative expression at each location (**Supplementary Fig. 4a**). An alternative binary visualisation mode highlights locations with nonzero expression of the selected gene, enhancing the visibility of gene activity (**Supplementary Fig. 4a**). Users can adjust the visualization granularity by modifying location size or filtering out low-expression locations, applying thresholds to streamline the display (**Supplementary Fig. 6e**). If SVGs were identified using the AW, these genes are highlighted with a turquoise-coloured result button for easy identification (**Supplementary Fig. 6b**).

Superimposition of other metadata, such as H&E staining or clustering results from Leiden or spatial clusters calculated in the AW, is facilitated for enhanced data interpretation (**Supplementary Fig. 6d**). When available, serial sections across a sample enable the uploading of a 3D model to improve spatial and anatomical orientation (**Supplementary Fig. 4c**), with various adjustments available to align the 3D model with the dataset.

The Visualiser’s side-by-side comparison feature allows visualization of two different gene expression profiles on adjacent slides or the same tissue section, enabling precise ROI delineation and ensuring identical analysis areas across slides. This feature can be toggled off to merge the expressions of both genes into a normalised, vector-based difference calculation at each location, highlighting significant expression differences between the two gene profiles (**Supplementary Fig. 4e-f**).

### Facilitating analysis of multi-slice Visium Datasets

For Visium datasets, VR-Omics overcomes two common analytical challenges: (1) facilitating concatenation of multiple Visium sections taken from the same tissue in 2D for joint analysis (**Supplementary Fig. 2e**) and (2) loading of multiple, serial sections for visualisation in 3D (**Supplementary Fig. 2f**). The first feature is enhanced through the introduction of a visually intuitive concatenation interface. Here, each sample is represented by its histologically stained image, allowing users to rearrange slides into their original configuration via a user-friendly drag-and-drop mechanism. Additionally, slides can be effortlessly rotated, enabling precise and flexible alignment for the accurate reconstruction of original tissue sections. Following alignment, users have the capability to specify processing parameters, including the filtering of cells, genes, and mitochondrial values, as well as adjusting the sensitivity of the clustering algorithm. These inputs initiate the seamless concatenation of slides, followed by subsequent analyses and cell type clustering, streamlining the entire process for optimal data integration and interpretation.

In the second scenario, VR-Omics offers the unique option to simultaneously load multiple Visium samples if they represent serial sections of the same tissue, to visualise the tissue in 3D (**Supplementary Fig. 2f**). For this, VR-Omics provides a straightforward alignment process consisting of selecting multiple Visium sample folders that have been processed and/or analysed with the AW (**Supplementary Fig. 2fi**). The selected datasets will be in unison, allowing the user to set the correct anatomical order from front to back of the slides. Next, all Haematoxylin and Eosin (H&E) images will be overlapped in a simple user interface that allows the user to rotate the slides individually, correcting misalignments of the samples on the capture areas (**Supplementary Fig. 2fii**). Furthermore, the user can set the distances between each slide in depth direction (**Supplementary Fig. 2fii**). Finally, the user can visualise the merged dataset rendered from the selected slides with the set distances and rotations as a 3D object (**Supplementary Fig. 2fiii**).

The virtual reconstruction offered through VR-Omics GUI allows datasets to be interpreted by biologists in a representation that is more faithful to the whole tissue. To provide training for prospective users, the AW and Visualiser capabilities can be explored by processing of the 3D demo data of the human developing heart [22]. When dealing with 3D datasets, the user can upload a 3D model of their tissue or organ of interest and align spatial data within it. In the case of the developing human heart [22], an overlay of the 3D model provides the user a more intuitive understanding of which clusters correspond to anatomical structures within the heart, allowing for easier selection of biologically relevant regions of interest (ROIs) across the depth of the dataset (Supplementary Fig. 4d).

### Processing patient samples and data analyses

Human ethics for cardiac tissue biobanking and human pluripotent stem cell research was approved by The Royal Children’s Hospital Melbourne Human Research Ethics Committee (approval numbers: HREC 38192). Informed consent was obtained from all human participants, and studies were conducted in accordance with the approved protocols and guidelines from the National Health and Medical Research Council of Australia. All samples were sequenced using the Visium FFPE Spatial Solution in conjunction with the Advanced Genomics Facility at the Walter and Eliza Hall Institute (WEHI). Samples of patient MCTHB302 were sequenced in the first batch, while the samples of MCTHB361 and of MCTHB362 were sequenced in a subsequent batch. Samples were sectioned as necessary to fit Visium capture areas (**Fig. 3,5 and Supplementary Fig. 8).** Outputs from both batches of Visium experiments were reanalysed through the SpaceRanger pipeline v2.0.1 [37] by the WEHI team. The outputs of the SpaceRanger pipeline for all samples were used as the inputs VR-Omics. This included using the Visualiser to align multiple 2D slices and generated a joint, co-planar data object (**Supplementary Video 1**), and Scanpy pre-processing including clustering using STAGATE. Filter parameters were established independently for each sample. Then markers for each cluster were used as the input for gene ontology enrichment analysis using Metascape [38] or Webgestalt [39].

These findings were then validated orthogonally, outside of the VR-Omics framework using a series of comparable tools requiring coding experience. Samples were pre-processed and filtered in Seurat then clustering for individual or joint objects for all patients was performed using the BayesSpace algorithm [40]. Gene ontology enrichment analysis was performed as outlined above.

## Availability of data and materials

The project home page providing VR-Omics software, tutorial, and demos is available under https://ramialison-lab.github.io/pages/vromics.html.

VR-Omics can also be accessed directly via https://figshare.com/articles/software/VR-Omics_Exploration_of_Spatial_Transcriptomics_data_in_3D/24503611. The source code can be accessed via https://github.com/Ramialison-Lab/VR-Omics

The paediatric cardiac rhabdomyomas patient Visium profiling data are available on NCBI GEO with accession number GSE252228. Third-party datasets used in this study: EGAS00001003996 [22]. Data for cross-platform comparison are available at https://figshare.com/s/78bcc6320cf3b2ee64af.

Custom R scripts for visualising Stitch3D data are available at https://figshare.com/s/0c9d9cc265b45b5d9441.

## Supplementary Information

### Supplementary Results

Additional results for cross-platform comparisons of mouse brain slides across multiple ST platforms are available. More information on multi-slice 3D data analysis using VR-Omics and orthogonal validation of up-regulated pathways in cRMs is also provided.

### Supplementary Methods

Additional information about the input data for ST platforms, the technical specifications of VR-Omics and its corresponding AW implementation, and the features of VR-Omics is provided. Furthermore, there is additional information on the analysis of cRMs using BayesSpace.

Supplementary Table 1: Benchmarking use case comparison and scoring between VR-Omics, STich3D, VR-Cardiomics, Loupe Browser, Xenium Desktop Browser, MERSCOPE Web-Vizualizer, Stereopy and SODB.

Supplementary Table 2: List of differentially expressed genes between the right ventricle and right atrium identified in the human embryonic heart at 6 dpc by VR-Omics.

Supplementary Table 3: Overview of all features supported by VR-Omics indicating for which SRT method they are available.

Supplementary Table 4: Feature comparison of VR-Omics with commonly used visualisation tools for each of the SRT methods supported by VR-Omics. The features described in this table show the overall capability of VR-Omics compared to the tools that are available to visualise and explore SRT data.

Supplementary Video: Overview of capabilities of VR-Omics.

## AUTHOR CONTRIBUTIONS

Conception and design: DB, NC, ERP, DAE, KK, HTN, FS, MR. Data acquisition, analysis and interpretation: DB, NC, HC-S, DM, BC, ATP, ERP, DAE, HTN, MR. Software development: DB, NC, DW, SJ-H, DG, JS, HTN. Manuscript draft: DB, NC. Manuscript editing and review: all authors.

## ACKNOWLEDGEMENTS

We thank the patient donors and their family for the valuable samples. We thank the members of the Ramialison’s, Schreiber’s, Porrello’s, and Elliott’s lab, especially Antonia Zech, Sebastian Bass-Stringer and Grzegorz Maciag for helpful discussions. We thank Celine Vivien for early staining and imaging of cRM samples that later informed experimental design. We thank the support of the WEHI Advanced Genomics Facility for technical advice on processing the patient samples. We thank Brett Kennedy and Patrick Marks (10x Genomics) and Jan Philipp Junker and Jeroen Bakkers for the sample data. We thank Yuen Chang and Jacquie Crawford and Francesca Bolk for support and assistance. We thank Geoff McDermott and Catherine King from 10x Genomics for their support and advice with the Visium platform. We thank Lisa Waylen and Maria Nucera for their helpful feedback on manuscript drafts.

---

**Supplementary Figure 1.** Top GO terms identified in Metascape when using the top differentially expressed genes (DEGs) between the right atrium and right ventricle using VR-Omics from the workflow to perform a 3D reconstruction of the human developing heart.

**Supplementary Figure 2.** Automated Workflow (AW). **a**, Data processing and filtering. Accepted SRT data types for analysis, processing and filtering using the AW include Visium, STOmics, MERFISH and Xenium. Analysis steps available to all SRT data types and menu with examples of available visualisations after data analysis. On the right, an example of heatmap of read counts generated for invasive ductal carcinoma breast tissue data generated with Visium from 10X Genomics [41]. **b**, Leiden or STAGATE clustering analysis and visualisation of clustering output using Visium data of the human cerebral cortex. **c**, Identification of SVGs is performed using SpatialDE for Visium data and by calculating Moran’s I statistic for MERFISH, STOmics and Xenium. **d**, Examples of output files generated by AW workflow with example data from coronal mouse kidney section generated by Visium. **e**, The user interface allows the multiple, co-planar 2D slices to be uploaded in tandem and aligned to form a single data object for processing. **f**, Multiple, serial Visium slides can be (**f_i_**) uploaded in tandem (**f_ii_**) so they can be combined into a 3D object (**f_iii_**), demonstrated here with data from the human embryonic heart at 6 dpc. The tissue images are used for reference.

**Supplementary Figure 3.** VR-Omics architecture. **a**, VR-Omics will accept as input sequencing and imaging-based SRT data of different technologies. **b**, The AW processes, filters and performing spatial analysis and producing output files. **c**, Data can be explored using VR-Omics as a 3D Visualiser in a virtual environment or desktop application as shown here with human embryonic heart at 6 dpc [22]. **d**, Output files, screenshots and video recordings are generated from the Visualiser.

**Supplementary Figure 4.** Features of the Visualiser. **a**, Gene search. **a_i_**, Gene search bar options for visualisation of selected gene expression patterns. **a_ii_**, Gene expression scale can be visualised with a colour map. The colour map uses a blue to red gradient, with lower expression levels indicated in blue and higher expression levels in red. The image represents the *EPCAM* gene expression in a breast tumour section (Xenium). **a_ii_**, Binary gene expression can be visualised. **b**, Visualisation of different dimensionality reduction methods, either projected back onto the tissue coordinates or as a UMAP or t-SNE pot. **c**, 3D model overlay. **c_i_,** Required inputs. 3D-model can be overlayed over single or multiple SRT slides in the Visualiser then aligned and orientated. **c_ii_,** Example of 3D model overlay displayed with data from the developing human heart. **c_iii_**, Accepted file formats. **c_iv_**, Available alignment movements. **d**, Selection of ROIs throughout slices and output. **d_i_**, User-defined selection of ROIs within 2D slide with option to select same ROI across serial 3D sections or to select up to 4 ROIs within an object. **d_ii_**, Visualisation of clusters in MERFISH adult mouse brain and zoomed individual clusters. **e**, Side-by-side comparison. **e_i_,** Allows comparison of two different gene profiles within identical copies of the same section for comparison of marker genes. **e_ii_**, Side-by-side comparison displayed using a Xenium breast tumour section^6.^ Expression of *EPCAM* is displayed in the left panel and expression of *ELF3* in the right panel. **f**, After side-by-side comparison has been selected, deselecting this option will ask the user to merge the slides. This creates a comparison of gene expression within the tissue section. Areas with more similar expression between the two genes will be displayed in cool colours, with areas of difference in warm colours, displayed here with the Visium human lymph node.

**Supplementary Figure 5.** Additional Visualiser features. **A**, Selection of screenshot function within the Visualiser for export of .png files, displayed here with 3D data from human embryonic heart at 6 dpc. **b**, Within the Visualiser, seamless zoom functionality allows the user to more closely inspect particular areas of the tissue, such as in the MERFISH adult mouse data. **c**, Interactive cluster visualisation within the Visualiser allows the user to visualise individual selected clusters across the tissue.

**Supplementary Figure 6.** Additional VR-Omics features. **a**, Example of format supported for custom data upload and example of processing and visualisation. Data from coronal mouse kidney section generated by Visium visualised here. **b**, SVG display. **b_i_,** Statistically significant SVGs will appear in the search blue highlighted in blue and a table containing additional information of all identified SVGs can be accessed in the Visualiser, displayed for the coronal mouse Visium brain slice. **b_ii_**, Example of additional information related to SVGs identified by Moran’s I statistic for Xenium and MERFISH data displayed for Xenium breast tumour section. **c**, Additional demo data availabe within VR-Omics for supported data generation platforms. **d**, Tissue image overlay between spots on slide and H&E stained image of the issue. The opacity of the images can be adjusted for maximise alignment of this data. **e**, After searching and selecting for a gene, a minimum expression threshold can be set to only display spots with a normalised expression above set threshold. Displayed here for Xenium breast tumour section. **e**_i_, scale of expression with cooler colours denoting lower expression and warmer colours denoting higher expression. Displayed here with gene *STC1*. **e**_ii_, Alternatively once the threshold has been set, expression of a gene such as *HARCR2* displayed here, can be displayed in a binary manner as being ‘on’ or ‘off’ above the threshold.

**Supplementary Figure 7.** Cross-platform comparison of data generated from a mouse brain coronal dataset, all visualised within the VR-Omics framework.

**Supplementary Figure 8.** Additional clustering data of cardiac rhabdomyomas. **a,** STAGATE labelled clusters using VR-Omics for analysis of MCHTB361 sample 2. **b,** STAGATE labelled clusters using VR-Omics for analysis of MCHTB361 sample 3**. c,** STAGATE labelled clusters using VR-Omics for analysis of MCHTB361 sample 1. **d**, BayesSpace joint clustering of multi-slice, co-planar MCHTB302 dataset used for orthogonal validation of results generated within VR-Omics. **e**, STAGATE labelled clusters using VR-Omics of multi-slice, co-planar MCHTB362 sample 2.

**Supplementary Figure 9:** GO analysis of cardiac rhabdomyomas using Metascape. **a**, GO results from MCHTB361 sample 2. **b**, GO results from MCHTB361 sample 3. **c**, GO results from MCHTB362 multi-slice, co-planar dataset sample 2.

