## Supplementary Table 1 for "Automated Integration of Multi-Slice Spatial Transcriptomics Data in 2D and 3D"

### **Supplementary Table 1:** Benchmarking use case comparison and scoring between VR-Omics, STich3D, VR-Cardiomics, Loupe Browser, Xenium Desktop Browser, MERSCOPE Web-Vizualizer, Stereopy and SODB.

| **Step** | **VR-Omics** | **STich3D** | **VR-Cardiomics** | **Loupe Browser** | **Xenium Desktop-Explorer** | **MERSCOPE Web Vizualizer** | **Stereopy** | **SODB** |
| --- | --- | --- | --- | --- | --- | --- | --- | --- |
| 1- Input Data | 1 | 1 | 0.5 | 1 | 1 | 1 | 1 | 1 |
| 2- Cluster the spots/locations | 1 | 1 | 1 | 1 | 1 | 0.5 | 1 | 1 |
| 3- Assemble the slices | 1 | 1 | 0 | 0 | 0 | 0 | 0 | 0 |
| 4- Manually select two ROIs | 1 | 0.5 | 1 | 0.5 | 0.5 | 0.5 | 0.5 | 0.5 |
| 5- Calculate the top DEGs between two ROIs | 1 | 0.5 | 1 | 0.5 | 0.5 | 0.5 | 0.5 | 0.5 |
| **Total Score** | **5** | **4** | **3.5** | **3** | **3** | **3** | **3** | **3** |
