## Supplementary Table 2 for "Automated Integration of Multi-Slice Spatial Transcriptomics Data in 2D and 3D"

**Supplementary Table 2:** List of differentially expressed genes between the Right Ventricle and Right Atrium identified in the human embryonic heart at 6 dpc^28^ by VR-Omics.

| **ENSEMBL ID** | **Gene NAME** | **logFC(Right Ventricle/Right Atrium)** | **P Value** | **Adjusted P Value** | **Average Expression** |
| --- | --- | --- | --- | --- | --- |
| ENSG00000111245 | MYL2 | 2.964946941 | 1.33E-137 | 1.38E-133 | 10.82222644 |
| ENSG00000092054 | MYH7 | 2.914745384 | 2.61E-152 | 5.38E-148 | 12.776794 |
| ENSG00000154556 | SORBS2 | 1.973076604 | 1.13E-61 | 7.78E-58 | 10.07464416 |
| ENSG00000134369 | NAV1 | 1.73851901 | 2.30E-54 | 1.19E-50 | 9.010788458 |
| ENSG00000160808 | MYL3 | 1.646547374 | 1.13E-40 | 4.66E-37 | 11.86476121 |
| ENSG00000155657 | TTN | 1.557131483 | 1.67E-27 | 2.87E-24 | 13.2862565 |
| ENSG00000213626 | LBH | 1.550151922 | 6.12E-40 | 2.11E-36 | 9.410438645 |
| ENSG00000038427 | VCAN | 1.505285595 | 1.64E-34 | 4.85E-31 | 9.257951681 |
| ENSG00000183023 | SLC8A1 | 1.472621156 | 6.24E-32 | 1.61E-28 | 10.62338056 |
| ENSG00000129991 | TNNI3 | 1.408416922 | 1.91E-28 | 3.95E-25 | 9.690107164 |
| ENSG00000159251 | ACTC1 | 1.331097373 | 5.28E-27 | 8.39E-24 | 12.147101 |
| ENSG00000159173 | TNNI1 | 1.319987129 | 4.97E-26 | 7.33E-23 | 11.15874395 |
| ENSG00000163399 | ATP1A1 | 1.133792727 | 1.37E-19 | 1.77E-16 | 9.420647966 |
| ENSG00000140416 | TPM1 | 1.10374865 | 1.25E-19 | 1.72E-16 | 12.89585187 |
| ENSG00000134571 | MYBPC3 | 1.094622511 | 9.38E-18 | 1.02E-14 | 10.33166835 |
| ENSG00000101605 | MYOM1 | 1.087395592 | 2.06E-17 | 2.12E-14 | 8.943975138 |
| ENSG00000114854 | TNNC1 | 1.056642352 | 2.90E-16 | 2.72E-13 | 11.57488477 |
| ENSG00000123416 | TUBA1B | 1.036389457 | 5.84E-19 | 7.09E-16 | 10.86619219 |
| ENSG00000065978 | YBX1 | 1.024028764 | 7.38E-16 | 6.10E-13 | 11.54710985 |
| ENSG00000198959 | TGM2 | 1.012374491 | 3.79E-15 | 2.90E-12 | 9.165397374 |
| ENSG00000170345 | FOS | -1.000342349 | 9.52E-13 | 4.92E-10 | 7.162486235 |
| ENSG00000185551 | NR2F2 | -1.081622198 | 2.09E-14 | 1.44E-11 | 7.238404227 |
| ENSG00000145730 | PAM | -1.667241703 | 6.66E-19 | 7.65E-16 | 8.350856287 |
| ENSG00000139219 | COL2A1 | -1.839595394 | 9.60E-28 | 1.80E-24 | 7.341104262 |
| ENSG00000197616 | MYH6 | -2.533851739 | 2.93E-30 | 6.74E-27 | 8.940247332 |
