## Supplementary Table 3 for "Automated Integration of Multi-Slice Spatial Transcriptomics Data in 2D and 3D"

**Supplementary Table 3:** Overview of all features supported by VR-Omics indicating for which SRT method they are available

|  |  | Visium | Xenium | MERFISH | STOmics | Tomo-Seq | Custom |
| --- | --- | --- | --- | --- | --- | --- | --- |
| **Automated Workflow (AW)** | **Filter dataset** | Yes | Yes | Yes | Yes | No | No |
|  | **Pre-process dataset** *(e.g., normalisation, log transformation, highly variable gene annotation)* | Yes | Yes | Yes | Yes | No | No |
|  | **Dimensionality Reduction:**   - *PCA* - *UMAP* - *t-SNE* | Yes | Yes | Yes | Yes | No | No |
|  | **Spatial analysis** *(e. g., Total_counts, N_genes_by_count, spatial co-occurrence, Ripley’s L function, neighbourhood enrichment analysis)* | Yes (partially) | Yes (partially) | Yes (partially) | Yes (partially) | No | No |
|  | **Clustering analysis** | Yes (Leiden + STAGATE) | Yes (Leiden) | Yes (Leiden) | Yes (Leiden) | No | No |
|  | **Spatially variable gene (SVG) analysis** | Yes (SpatialDE) | Yes (Moran’s I) | Yes (Moran’s I) | No | No | No |
|  | **Generation of output plots** | Yes | Yes | Yes | Yes | No | No |
|  | **Results as CSV** | Yes | Yes | Yes | Yes | No | No |
|  | **Process + analyse a joint dataset from multiple 2D experiments** | Yes | No | No | No | No | No |
| **Visualiser** | **VR visualisation** | Yes | Yes | Yes | Yes | Yes | Yes |
|  | **Desktop 3D visualisation** | Yes | Yes | Yes | Yes | Yes | Yes |
|  | **GUI to join multiple 2D slides in 3D dataset** | Yes | No | No | No | No | No |
|  | **GUI to join multiple 2D slides in 2D dataset** | Yes | No | No | No | No | No |
|  | **Demo data available** | Yes | Yes | Yes | Yes | Yes | No |
|  | **Selection of ROIs** | Yes | Yes | Yes | Yes | No | Yes |
|  | **Side-by-side comparison** | Yes | Yes | Yes | Yes | No | Yes |
|  | **Comparison of two gene profiles** | Yes | Yes | Yes | Yes | No | Yes |
|  | **Single location interaction** | Yes | Yes | Yes | Yes | No | Yes |
|  | **Heatmap visualisation of gene profile** | Yes | Yes | Yes | Yes | Yes | Yes |
|  | **Binary visualisation of gene profile** | Yes | Yes | Yes | Yes | No | Yes |
|  | **Visualisation of cluster information** | Yes | Yes | Yes | Yes | Yes | Yes (if provided) |
|  | **Minimum threshold for gene expression** | Yes | Yes | Yes | Yes | Yes | Yes |
|  | **SVG highlighted in Visualiser** | Yes | Yes | Yes | No | No | No |
|  | **3D model overlay** | Yes | Yes | Yes | Yes | Yes | Yes |
|  | **Image overlay (e. g. H&E Stain)** | Yes | No | No | No | No | Yes (if provided) |
|  | **Customisable environment** *(e.g., Location symbol, Colour gradient, Dark/Light mode)* | Yes | Yes | Yes | Yes | Yes | Yes |
|  | **Dimensionality reduction (UMAP or t-SNE) visualisation** | Yes | Yes | Yes | Yes | No | No |
|  | **Visualisation of additional values such as** *(e. g., genes_by_count, total_counts, mt_counts)* | Yes | Yes (partially) | Yes (partially) | No | No | No |
| **Detailed documentation for AW and Visualiser available** | | Yes | Yes | Yes | Yes | Yes | Yes |
