## Supplementary Table 4 for "Automated Integration of Multi-Slice Spatial Transcriptomics Data in 2D and 3D"

**Supplementary Table 4:** **Comparison of VR-Omics with commonly used visualisation tools for each of the SRT methods supported by VR-Omics.** The features described in this table show the overall capability of VR-Omics compared to the tools that are available to visualise and explore SRT data.

|  | **VR-Omics** | **Loupe Browser (Visium)** | **Xenium Desktop-Explorer**  **(Xenium)** | **MERSCOPE Web-Vizualizer**  **(MERFISH)** | **SODB** | **VT3D** | **Spateo** | **Stitch3D** | **Spacemake** | **StereoMap** |
| --- | --- | --- | --- | --- | --- | --- | --- | --- | --- | --- |
| **Analysis options** | | | | | |  |  |  |  |  |
| *3D object overlay* | Yes | No | No | No | No | Yes | Yes | Yes (computational skills required) | No | No |
| *Joint analysis of multiple 2D slides* | Yes | No | No | No | No | No | No | No | No | No |
| *3D SRT dataset alignment and exploration of multiple slides* | Yes | No | No | No | No | No | Yes | Yes | Yes | No |
| *Analysis and display of spatially variable genes* | Yes | Partially (display of pre-annotated SVGs) | No | No | Partially (display of pre-annotated SVGs) | No | Yes (computational skills required) | No | Yes (computational skills required) | No |
| *Customised sub selection of ROIs* | Yes | Yes | Yes | No | Yes | No | Yes | No | No | Yes |
| *Side-by-Side comparison of two SRT slides* | Yes | No | No | No | Yes | No | No | No | No | No |
| *Support multiple SRT technologies* | Yes | No | No | No | Yes | Yes | Yes | Yes | Yes | No |
| *Visualisation of cell boundaries (segmentation)* | No | No | Yes | Yes | No | No | Yes | No | Yes | No |
| **Import/Export options** | | | | | |  |  |  |  |  |
| *Import SRT data* | Yes | Yes | Yes | Yes | Partially (computational skills required) | Partially (computational skills required) | Partially (computational skills required) | Partially (computational skills required) | Partially (computational skills required) | Yes |
| *Supporting data as provided by manufacturer* | Yes | Yes | Yes | No | No | No | Yes | Yes | Yes | Yes |
| *Import SRT data with x, y and z coordinates (Custom data)* | Yes | No | No | No | No | Yes | Partially (computational skills required) | Partially (computational skills required) | No | No |
| *Cross platform support of multiple SRT platforms* | Yes | No | No | No | Yes | Yes | Yes | Yes | Yes | No |
| *Export options*   - *Screenshot/ Figure plots* - *Files* - *ROI selections* - *Video/mp4* | Yes  Yes  Yes  Yes | Yes  Yes  Yes  No | Yes  No  Yes  No | Yes  Yes  Yes  No | Yes  Yes  No  No | No  No  No  No | Partially (computational skills required) | Partially (computational skills required) | Partially (computational skills required) | Yes  Partially (Status file only)  No  No |
| **Performance** | | | | | |  |  |  |  |  |
| *GPU instancing to improve performance for large datasets* | Yes | No | No | No | No | No | No | No | No | No |
| *Integrated Automated Workflow for data mining/processing* | Yes | No | No | No | No | No | No | No | Partially (computational skills required) | No |
| **User-friendliness** | | | | | |  |  |  |  |  |
| *Documentation*   - *Video tutorial* - *Pictures step by step* | Yes  Yes | No  Yes | No  Yes | Yes  No | Yes  Yes | No  No | Yes  Yes | No  Yes | No  Yes | No  No |
| *Interactive non-computational User Interface (GUI)* | Yes | Yes | Yes | Yes | Partially | Partially | No (Only ROI selection) | No (Only interactive plots, computational skills required) | No | Yes |
| *Local desktop installation* | Yes | Yes | Yes | Yes | Partially (local installation required for analysis) | Yes  (Computational skills required) | Yes  (Computational skills required) | Yes  (Computational skills required) | Yes  (Computational skills required) | Yes |
| *Online access* | No | No | Yes | Yes | Yes | Yes (AtlasBrowser) | No | No | No | No |
| *Open source* | Yes | No | No | No | Yes | Yes | Yes | Yes | Yes | No |
| *VR mode* | Yes | No | No | No | No | No | No | No | No | No |
